# Allometric multi-scaling of weight-for-height relation in children and adolescents: Revisiting the theoretical basis of body mass index of thinness and obesity assessment

**DOI:** 10.1101/2024.03.13.584792

**Authors:** Hitomi Ogata, Sayaka Nose-Ogura, Narumi Nagai, Momoko Kayaba, Yosuke Isoyama, João Kruse, van Seleznov, Miki Kaneko, Taiki Shigematsu, Ken Kiyono

**Affiliations:** Graduate School of Humanities and Social Sciences, Hiroshima University, Hiroshima 739-8521, Japan; JAPAN High Performance Sport Center, Japan Institute Sports Sciences, Department of Sports Medicine and Research, Tokyo 115-0056,Japan; Department of Obstetrics and Gynecology, University of Tokyo Hospital, Tokyo 113-8655, Japan; School of Human Science and Environment, University of Hyogo, Himeji, 670-0092, Japan; Faculty of Medicine, University of Tsukuba, Tsukuba, 305-8577, Japan; Graduate School of Engineering Science, Osaka University, Osaka 560-8531, Japan

## Abstract

The body mass index (BMI), defined as weight in kilograms divided by height in meters squared, has been widely used to assess thinness and obesity in all age groups, including children and adolescents. However, the validity and utility of BMI as a reliable measure of nutritional health have been questioned. This study discusses the mathematical conditions that support the validity of BMI based on population statistics. Here, we propose a condition defined as allometric uni-scaling to ensure the validity of BMI as an objective height-adjusted measure. Any given centile curve, including the median curve, in a weight-for-height distribution should be approximated using power-law functions with the same scaling exponent. In contrast, when the scaling exponent varies depending on the position of the centile curve, it is called allometric multi-scaling. By introducing a method for testing these scaling properties using quantile regression, we analyzed a large-scale Japanese database that included 7,863,520 children aged 5-17 years. We demonstrated the remarkable multi-scaling properties at ages 5-13 years for males and 5-11 years for females, and the convergence to uni-scaling with a scaling exponent close to 2 as they approached 17 years of age for both sexes. We confirmed that conventional BMI is appropriate as an objective height-adjusted mass measure at least 17 years of age, close to adulthood, for both males and females. However, the validity of BMI could not be confirmed in younger age groups. Our findings indicate that the growth of children’s weight-for-height relation is much more complex than previously assumed. Therefore, a single BMI-type formula cannot be used to assess thinness and obesity in children and adolescents.

## Introduction

The body mass index (BMI), defined as weight in kilograms divided by height in meters squared, has been widely used to assess both body thinness and fat in all age groups, including children and adolescents [1–4]. In practice, the World Health Organization (WHO), medical doctors, nutritionists, and elementary schools use BMI to screen for thinness, underweight, overweight, and obesity [5, 6]. In recent years, however, negative opinions have emerged regarding the validity of BMI, including that BMI does not account for differences across race/ethnic groups, sex, gender, and age span; it is an inaccurate measure of body fat content and does not consider muscle mass, bone density, and overall body composition; and increased BMI is not always associated with increased cardiovascular and mortality risks. [7, 8].

As BMI may have limitations in its application and may not be a versatile measure for every purpose, it is crucial to establish scientific methods to verify its basis and validity. BMI-related issues can be categorized as follows (see Fig. 1): (1) basis for the BMI formula, (2) validity of BMI as an objective measure of thinness and fatness in a population with different heights, and (3) validity of BMI cutoff values for screening nutritional status. Answers to these problems could be provided by several scientific approaches, such as a physiological approach based on body composition measurements such as body fat percentage, an epidemiological approach based on disease prevalence, and a population statistical approach based on a statistical analysis of a large dataset consisting of height and weight [9–11]. This study primarily deals with the population statistical approach (hereafter referred to as the population-based approach).

**Fig 1.**
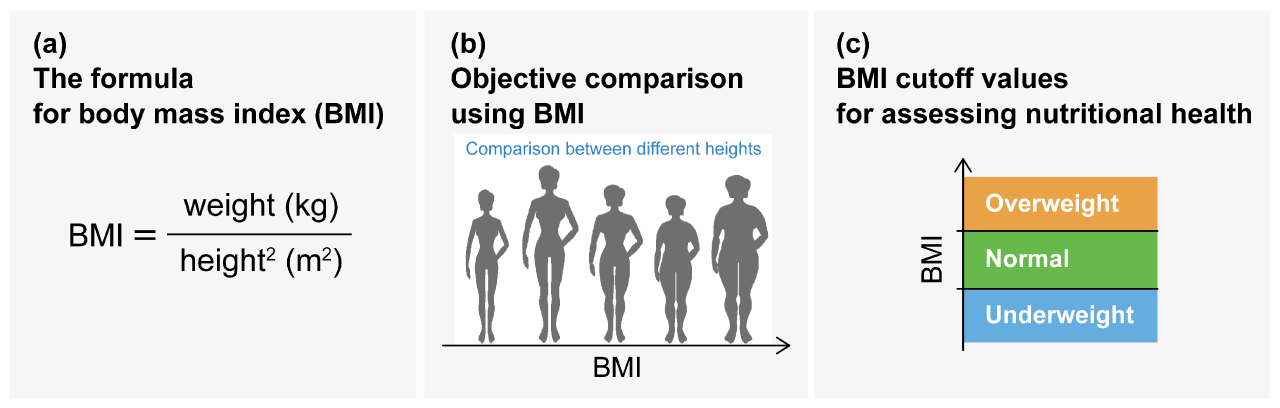
Classes of BMI validity issues. (a) The basis for the BMI formula: How is this formula derived? (b) Validity of BMI as an objective height-adjusted measure in a population with differences in height: Is BMI a height-independent measure? (c) Validity of BMI cutoff values: How do we determine the cutoff value?

To date, population-based approaches have mainly offered a foundation supporting BMI, although it is becoming physiologically evident that BMI is a poor indicator of body fat percentage [6, 12–14]. In the 1960s and the 1970s, in addition to BMI, formulas for (weight)/(height)^*α*^, such as the body build index (weight divided by height: *α* = 1) and the Rohrer index or inverse Ponderal index (weight divided by height cubed: *α* = 3), were also studied [15–19]. Remarkably, in 1835, Lambert Adolph Jacque Quetelet noted that the relationship between body weight and height in normal young adults was least affected by height when the ratio of weight to height squared was used, rather than the ratio of weight to height or weight to height cubed [12]. In 1972, this was re-validated by Keys et al. [12], who demonstrated the significance of BMI as the index with the most negligible correlation with height. In 1993, the WHO Health Organization assembled BMI cutoff values and nutritional status classifications, which are widely utilized today [20]. In early studies, BMI was validated based on the uncorrelated criterion that the correlation between BMI and height was negligibly weak [12, 21]. However, this study shows that the uncorrelated criterion must be revised as the basis for supporting an objective height-adjusted mass measure.

In mathematical biology and physiology, an allometric scaling relationship in the weight-for-height distribution provides the basis for the BMI formula [22, 23]. Allometry is the study of the relationship between the size of an organism and its physiology, morphology, and life history [24]. In most living organisms, body-size relationships are described by a power-law function called allometric scaling, *y* = *Cx*^*∝*^, where *x* is the organism’s body size, *y* is a biological attribute (trait), and *∝* and *C* are statistically estimated scaling exponents and constants, respectively. Based on allometric scaling, variations in body mass among individuals or species have been used to predict traits, such as metabolic rate, survival probability, and fecundity [25, 26]. The defining formula for BMI comes from a scaling law, *w* = *Ch*^*∝*^, where *w* and *h* are the weight and height of the general population, respectively. It is empirically known that the resting energy expenditure is proportional to *w*^3*/*4^ in a wide range of animals, such as unicellular organisms, metazoans, and thermostat animals (known as Kleiber’s law or 3*/*4-power law scaling) [24, 27]. Based on this fact and the assumption that energy expenditure is proportional to the body surface area to maintain a constant body temperature [28], we can mathematically infer that a human’s weight could be proportional to *h*^2^ [29] (see Results and Discussion for details).

Early studies, including relatively large samples (thousands of cases), also demonstrated that the scaling exponent in several different racial/ethnic adult groups is close to 2 [30, 31]. Based on the conventional allometric framework, we assume that *w/h*^*∝*^ provides a normalized weight index for height when *w* = *Ch*^*∝*^ holds true. Therefore, the empirical fact that *∝* = 2 provides the basis for the BMI formula *w/h*^2^. This fact provided the basis for the BMI formula (Fig. 1(a)) and does not provide any evidence for other BMI-related issues as shown in Figs. 1(b) and 1(c). Interestingly, Cole (1986) showed that the scaling exponent *α* changes with age, with *∝* = 2 in infancy, rising to 3 in adolescence, and falling back to 2 in adulthood [21, 31, 32]. Thus, the findings suggest that modifying the BMI formula using age-dependent *∝* is preferable [33].

To assess thinness, underweight, overweight, and obesity, BMI cutoff values were defined as shown by the solid lines in Fig. 2(b)). In the population-based approach, a BMI cutoff value was determined as a specified centile (or z-score) point of the BMI distribution consisting mostly of healthy people, rather than a cutoff value to classify disease and healthy groups. A centile (also called a percentile or quantile) is a value that represents a percentage position on a list of observed data sorted in increasing order. For instance, if we had weight data for 101 individuals and sorted them in increasing order, the minimum weight is the value of the 0th centile, the maximum weight is the value of the 100th centile, and the 3rd, 11th, 51st, 91st, and 99th ranked weights in order of the smallest to the largest are in the 2nd, 10th, 50th, 90th, and 98th centiles, respectively.

**Fig 2.**
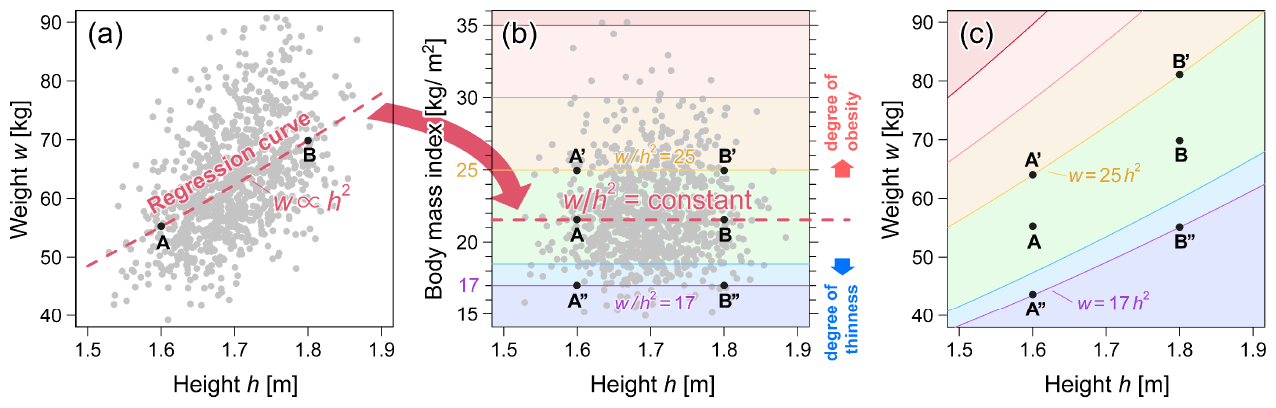
Relationship between weight-for-height and BMI-for-height distributions. (a) Example of weight-for-height distribution. The weight-for-height relationship (the median curve of the weight) can be empirically approximated by a power law of *w ∝ h*^2^, where *∝* denotes a proportional relation. (b) The BMI-for-height distribution converted from (a). The horizontal solid lines represent the cutoff values for thinness, underweight, overweight, and obesity from bottom to top. (c) Weight-for-height cutoff curves correspond to the BMI cutoffs shown in (b).

Notably, before discussing the validity of the BMI cutoff values, we should test the validity of BMI as an objective height-adjusted measure allowing comparison of populations with different heights. As illustrated in Fig. 2, and points A and B in Fig. 2(a), the two different weights with different heights on the regression curve described by *w* = *Ch*^2^ have the same BMI values as in Fig. 2(b). In this case, the regression curve corresponds to the median (50th centile) curve of the weight in the weight-for-height distribution. In the thinness and obesity assessment using BMI, points A and B, A’ and B’, and A” and B” in Fig. 2(b), were assumed to have a similar body build. It would be statistically sound to interpret that each pair of A and B, A’ and B’, and A” and B” pass through the same centile (quantile) curves of weight in the weight-for-height distribution, as shown in Fig. 2(c)). That is, a BMI cutoff value in Fig. 2(b) correspond to the specified centile curve in Fig. 2(c)). In other words, individuals with the same BMI were at the same centile (centile) in each height group. However, it is not apparent that this assumption holds true. If this assumption was not fulfilled, the validity of BMI as an objective height-adjusted measure would be rejected.

The main purpose of this study is to investigate this problem, which has never been pointed out or examined. In our population-based approach, the validation of BMI was tested based on whether any given centile curve of the weight-for-height distribution exhibited allometric scaling with the same scaling exponent, and not just a median curve. When all centile curves have the same scaling exponent, we refer to this type of allometric scaling as uni-scaling, which provides the basis for the validity of BMI. In contrast, when each centile curve exhibits allometric scaling with a different exponent, it is referred to as multi-scaling. In conventional allometry, uni-scaling has been implicitly assumed and multiple scales have not been hypothesized. In this study, we propose a method to assess the uni- and multi-scaling properties of weight-for-height distributions using quantile regression. By analyzing a large-scale database including 7,863,520 children aged 5-17 years, we demonstrated the existence of multi-scaling properties of weight-for-height distributions at ages 5-13 years for males and 5-11 years for females and the convergence to uni-scaling with a scaling exponent close to 2 as they approach 17 years of age for both sexes. Our findings indicate that although conventional BMI can be seen as a well-grounded measure, at least statistically, for young adulthood, a single index such as BMI cannot be used to assess both thinness and obesity in children and adolescents. In the second part of this paper, we discuss a method for assessing thinness in children and adolescents using the extended BMI instead of the conventional BMI.

## Extended allometric analysis: uni- and multi-scaling

Before describing the methods and results of our study, we illustrate the key ideas of our analysis and the new concepts in allometric scaling. Conventional allometric analysis implicitly assumes that the relationship between two variables in our study, weight *w* and height *h*, can be fully characterized by a single scaling exponent *∝*. In this situation, any given centile curve of the weight in the weight-for-height distribution can be approximated using a power-law function *w*(*h*) = *Ch*^*∝*^, with the same scaling exponent *∝*. To explicitly test this property, we fit the power-law function to *q*th centile curve, as shown by the solid line in Fig. 3 using the quantile regression method [34]. In this analysis, the scaling exponent *∝*(*q*) may vary depending on *q*. We refer to this analysis as extended allometric analysis.

**Fig 3.**
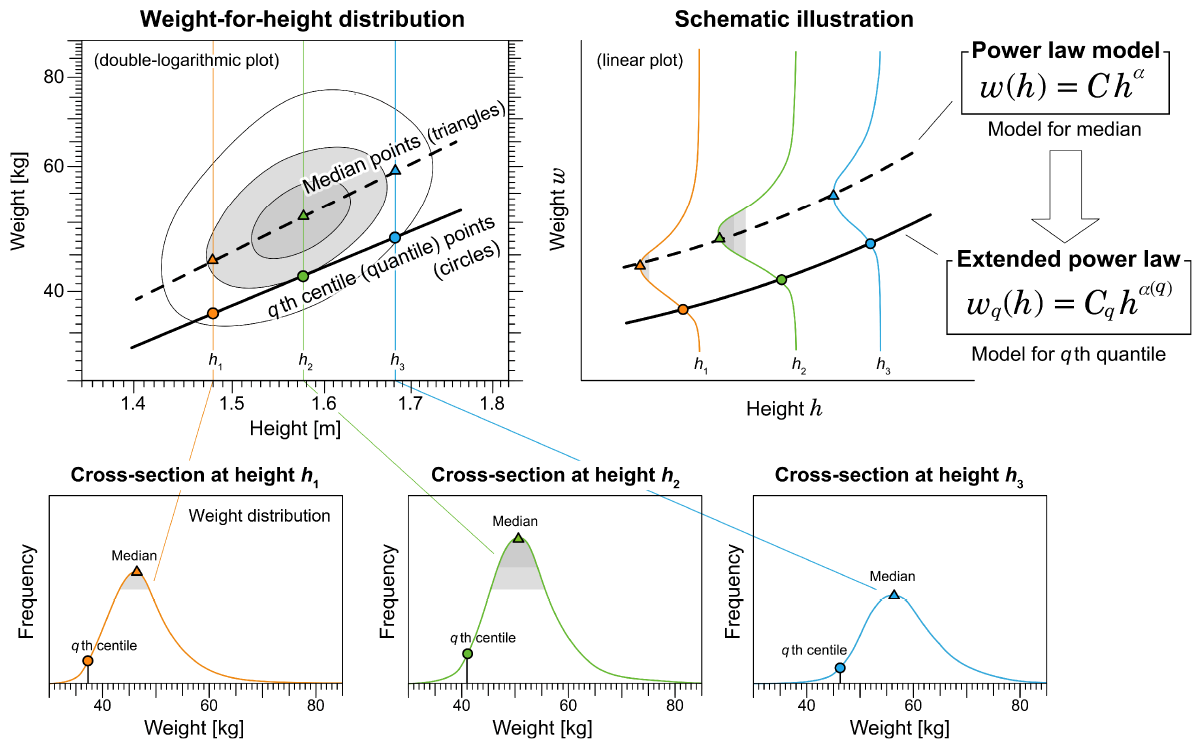
Schematic illustration of *q*th quantile (centile) regression. In the extended allometric analysis, the *q*th centile (quantile) curve of a weight-for-height distribution is fitted by a power-law function *w*_*q*_(*h*) = *C*_*q*_ = *h*^*∝*(*q*)^.

If the estimated exponents *{∝*(*q*)} are independent of *q* as illustrated in Fig. 4(a), we refer to this property as uni-scaling. In this case, BMI centile curves were constant and independent of height, as illustrated by the horizontal parallel lines in Fig. 4(c). By contrast, if the estimated exponents *{α*(*q*)*}* depend on *q* as illustrated in Fig. 4(b). We call this property multi-scaling. In this case, BMI centile curves depended on height, as illustrated in Fig. 4(d). The height dependence of BMI indicates that BMI is inappropriate as an objective and unified measure of body thinness and fat. During the uniscaling, as shown in Fig. 4(c), two points A and B with the same BMI are on the same 90th centile curve, which guarantees a statistical interpretation of BMI. In contrast, when multi-scaling, as illustrated in Fig. 4(c), two points A and B with the same BMI are on different centile curves, which means that BMI is difficult to interpret, at least statistically. As illustrated in Fig. 5, the uni-scaling condition can provide an objective interpretation of BMI, and the comparison of BMI corresponds to a comparison of centile positions of weight between populations of different heights.

**Fig 4.**
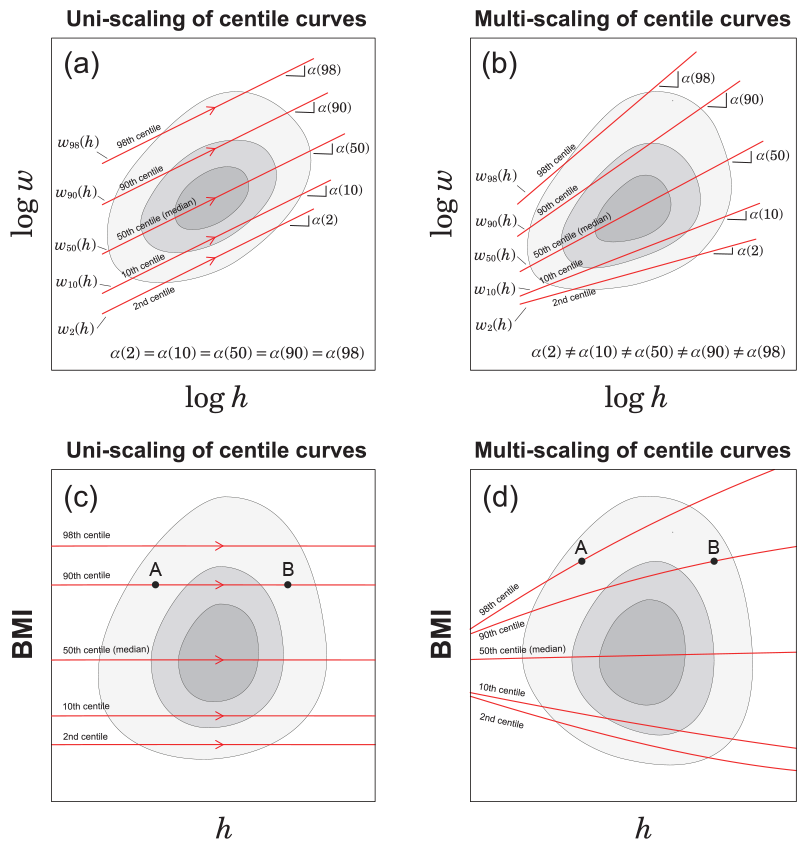
Schematic illustration of uni- and multi-scaling. (a) Uni-scaling in weight-for-height distribution. In a double-logarithmic plot, all centile curves are parallel straight lines. (b) Multi-scaling in weight-for-height distribution. In a double-logarithmic plot, slopes of the centile curves vary with position. (c) Uni-scaling centile curves in BMI-for-height distribution. When uni-scaling, two points with the same BMI are on the same 90th centile curve, A and B. (d) Multi-scaling centile curves in BMI-for-height distribution. When multi-scaling, two points with the same BMI are on different centile curves, A and B.

**Fig 5.**
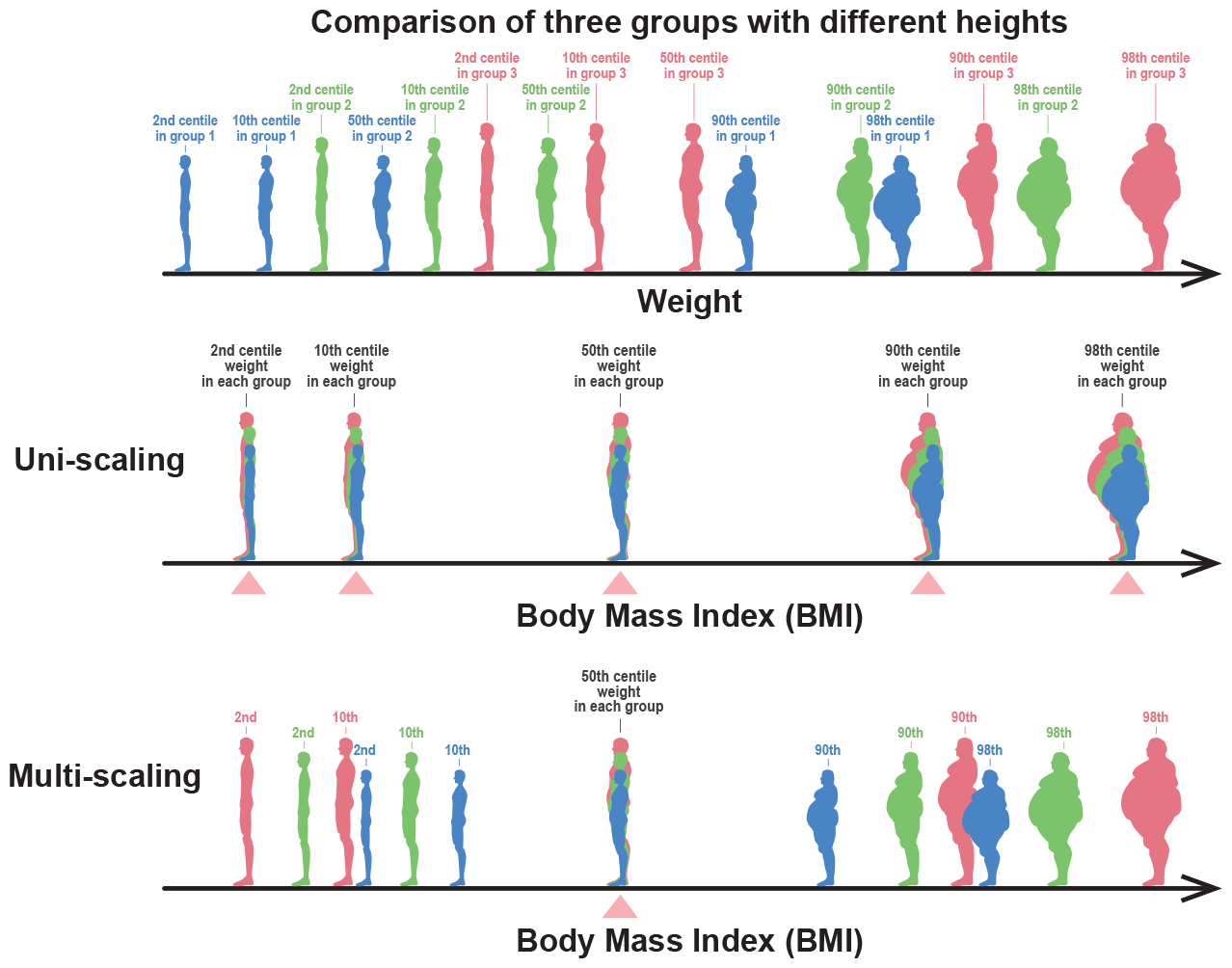
Meaning of the uni-scaling criterion. The upper figure schematically depicts a body weight distribution of people with 2nd, 10th, 50th, 90th, and 98th centile weights from the three different height groups (red, green, and blue represent the tall, medium, and short height groups, respectively). If the uni-scaling condition of the weight-for-height distribution is fulfilled, the BMI values place the weight centile points of each group in the same position (indicated by triangles), as depicted in the middle row. Thus, the comparison of BMI corresponds to a comparison of weight centile positions adjusted by heights. In contrast, if multi-scaling, the BMI cannot make the lower and higher centile positions comparable, although it makes the 50th centile (median) weight at the same position, as depicted in the bottom row.

## Data and method

We analyzed a large-scale national dataset including the age, sex, height, and weight of 7,863,520 children aged 5-17 years between 2008 and 2019, wherein 2.72% of the samples, including any missing values, were excluded from a total of 8,083,466 samples. The number of samples for each sex and age group is summarized in Table 1. To clarify the physical development of children in schools, data were collected from School Health Statistics Research conducted by the Ministry of Education, Culture, Sports, Science, and Technology in Japan. This national sample survey employed a stratified random sampling method wherein schools in each prefecture were stratified by the number of enrolled students to ensure coverage of small to large schools (in other words, to reduce the bias associated with the population size of municipalities); subsequently, within each school-size stratum, the surveyed schools were simply randomly selected. Finally, the children surveyed were selected using a systematic sampling method based on age and sex. The survey is conducted annually from April to June. However, after 2020 when the coronavirus disease 2019 pandemic occurred, the survey period could not be scheduled from April to June and was conducted year-round. Therefore, data obtained after 2020 were not used in our analysis.

**Table 1.** Number of samples for each age and sex after removing samples with missing values.

| Age<br>[years] | Number of samples |  | [years] | Male | Female |
| --- | --- | --- | --- | --- | --- |
|  | Male | Female |  |  |  |
| 5 | 358,375 | 355,134 | 12 | 427,950 | 428,705 |
| 6 | 260,536 | 260,090 | 13 | 428,295 | 429,324 |
| 7 | 261,178 | 260,666 | 14 | 429,119 | 429,896 |
| 8 | 261,374 | 260,903 | 15 | 239,003 | 241,696 |
| 9 | 261,852 | 261,426 | 16 | 238,935 | 241,314 |
| 10 | 261,898 | 261,643 | 17 | 238,851 | 240,958 |
| 11 | 262,405 | 261,994 | Subtotal | 3,929,771 | 3,933,749 |
|  |  |  | Total | 7,863,520 |  |

### Smoothed bootstrap

In the dataset, height and weight are recorded as integers in centimeters (cm) and kilograms (kg), respectively. We applied the smoothed bootstrap technique to smoothen the integer-valued discrete distributions of the original data and improve statistical confidence [35]. In this smoothed bootstrap technique, the rows in the dataset are randomly extracted by replacement to obtain samples of size *N* from an initial dataset of size *N* . Subsequently, as illustrated in Fig. 6, random noise from a bivariate Gaussian kernel density was added to each pair of heights and weights in the extracted row. The smoothing bandwidth is estimated using a rule-of-thumb bandwidth selector for bivariate Gaussian kernels [36]. In this study, the bootstrap technique was repeated 200 times and a statistical estimate was calculated for each smoothed bootstrap replication. In the following, we present the median values of all replicates as estimates.

**Fig 6.**
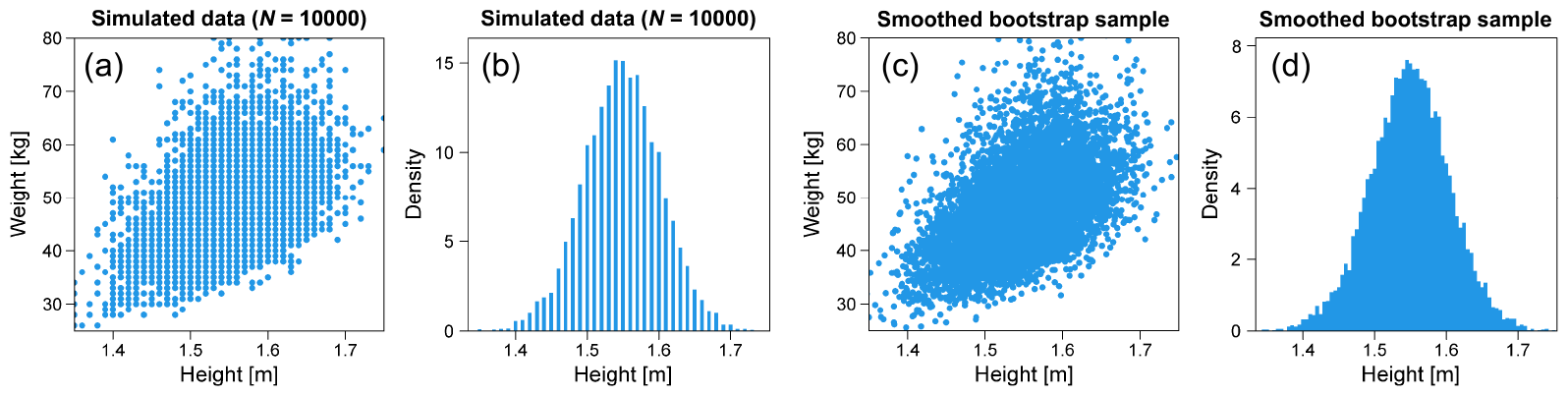
Smoothed bootstrap by adding noise. (a) Sample plot of integer height and weight data. (b) Histogram of integer height data. (c) Sample plot of height and weight data after adding noise. (d) Histogram of height data after adding noise.

### Extended allometric scaling for any given centile of weight-for-height distribution

We extended the allometric scaling to any given centiles of the weight-for-height distribution for each sex and age, as shown in Fig. 3. The *q*th centile of a continuous random variable *X* is defined as the value *x* such that Prob (*X ≤ x*) = *q/*100. Our study considers the conditional centile of the weight-given height and assumes that it follows an extended power-law model.

The extended power-law model is described as

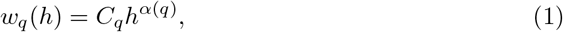

where *w*_*q*_(*h*) represents the *q*th centile body weight in kilograms (kg) given *h, h* represents height in meters (m), and the model parameters *α*(*q*) and *C*_*q*_ are the scaling exponent and proportionality constant, respectively. We employ a quantile regression approach to estimate the model parameters [34].

### Quantile regression

A quantile regression estimates the conditional quantiles (centiles) of a response variable [34]. For instance, in a quantile regression using an allometric model, we minimize the sum of the weighted absolute residuals:

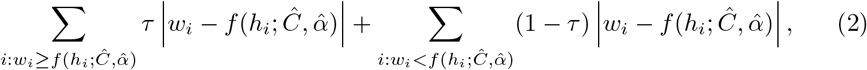

where weight *w*_*i*_ is the response variable, *i* is the index of an individual, *f* (*h*_*i*_; *C, α*) = *Ch*^*α*^ is the allometric model, *Ĉ* and 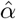 are the estimated values of the model parameters, and *τ* is set to *q/*100 when *q*th quantile regression is applied. To identify the model parameters, we used a simplex search algorithm [37].

## Results and Discussion

### Extended allometric analysis results

Figure 7 shows the results of the extended allometric analysis for 17-year males (a), females (b), and 8-year males (c) and females (d). As indicated by the solid lines in Figs. 7(a) and 7(b), the regression lines of the 2nd, 10th, 50th, 90th, and 98th centiles for 17-year-old males and females demonstrate almost uni-scaling, with a scaling exponent close to 2. In contrast, as indicated by the lines in Figs. 7(c) and 7(d), the regression lines of 8-year males and females demonstrate apparent multi-scaling properties.

**Fig 7.**
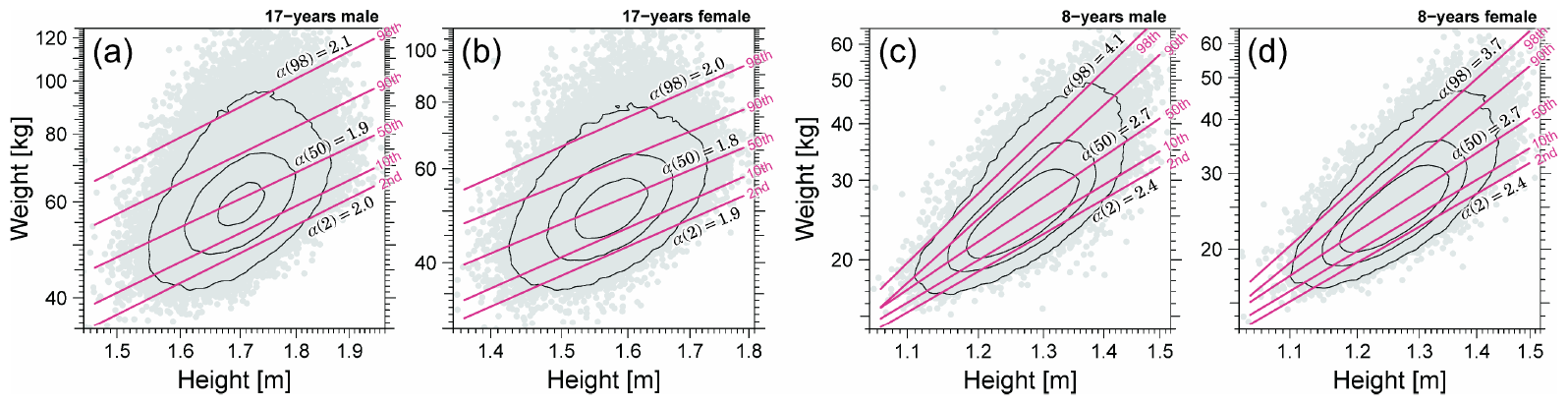
Extended allometric analysis results. (a) 17-year males. (b) 17-year females. (c) 8-year males. (d) 8-year males. Solid lines show the quantile regression results for 2nd, 10th, 50th, 90th, and 98th centiles in a double logarithmic scale. Estimated scaling exponents (slopes of the plots), α(2), α(50), and α(98), are shown in each panel.

Figure 8 shows the age dependence of scaling exponent *α*(*q*) for *q* = 2, 10, 50, 90, and 98. As shown in Figs. 8(a) and (b), although all scaling exponents *{α*(*q*)*}* converged to the same value close to 2 as the age of both males and females approached 17, multi-scaling with different scaling exponents *{α*(*q*)*}* was observed at younger ages. As indicated by the green lines in Figs. 8(a) and (b), the age dependence of α(50) corresponding to the conventional allometric scaling exponent varies with α(50) = 2 around 5 years, rising to 3 around 10 years, and then falling back to 2 in adulthood, as reported by Cole. However, the scaling exponents of higher and lower centiles, such as α(2) (light blue line) and α(98) (red line), showed different values from α(50) at ages 5-13 years for males and 5-11 years for females. Our findings would raise two issues regarding the use of conventional BMI in children. One is the BMI formula and the other is the objectivity of BMI even after modifying the BMI formula based on the estimated scaling exponent *α*(50). Before discussing these issues in detail, we explain in what sense BMI is a good objective measure for thinness and obesity assessment, and emphasize that BMI for adults is valid in the sense of population statistics.

**Fig 8.**
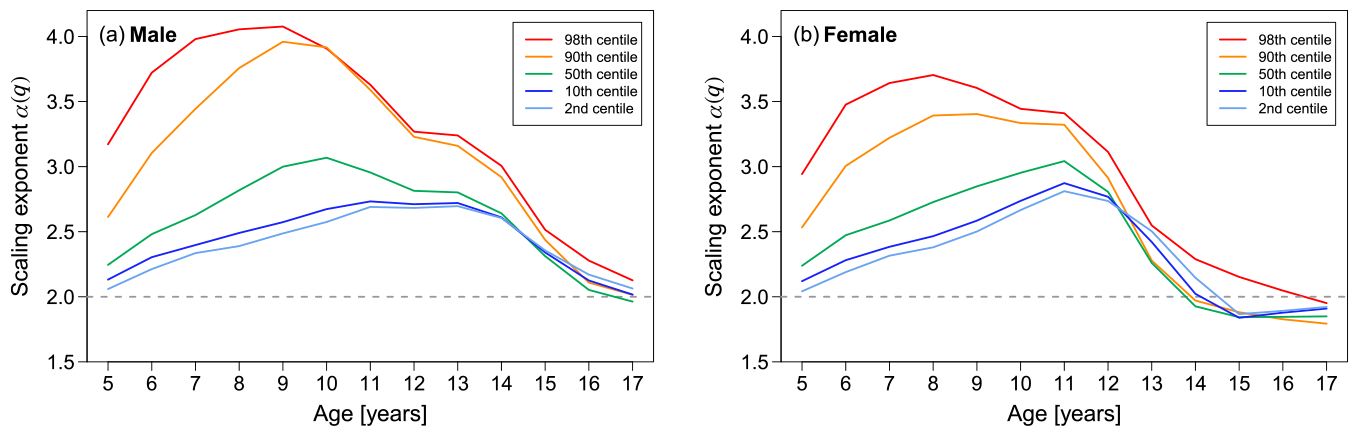
Age dependence of scaling exponents. The estimated scaling exponents of the 2nd, 10th, 50th, 90th, and 98th centiles for the weight-for-height distribution are plotted from bottom to top. (a) Male. (b) Female. If the five lines have almost the same value, it indicates uni-scaling; if the values are different, it indicates multi-scaling.

### Why use BMI? — Importance of uni-scaling criterion —

To explain the validity of BMI for adults, we compared the structures of the weight-for-height relationship (Fig. 9(a)) and BMI-for-height relationship (Fig. 9(b)) in 17-year females as close to adulthood. In Fig. 9(a), the 2nd, 50th, and 98th quantile regression curves assuming Eq. (1) to the weight-for-height data (gray points) are indicated by solid and dashed lines. In addition to the quantile regression curves, the estimated 2nd, 50th, and 98th centile points of the weight distribution in each stratum of height are plotted as circles, triangles, and diamonds, respectively, in Fig. 9(a), which is consistent with the corresponding quantile regression curves. As shown in Fig. 9(a), all the regression lines appear parallel for the 2nd, 50th, and 98th centiles of the weight-for-height distribution, indicating uni-scaling. If we define the standardized BMI as *w/h*^*α*(50)^ [kg/m^*α*(50)^] with *α*(50) = 1.85, the 2nd, 50th, and 98th centile levels of the BMI-for-height distribution would become constant and independent of height. Therefore, as illustrated in Fig. 5(middle row), the uni-scaling property guarantees to compare centile positions between different heights objectively.

**Fig 9.**
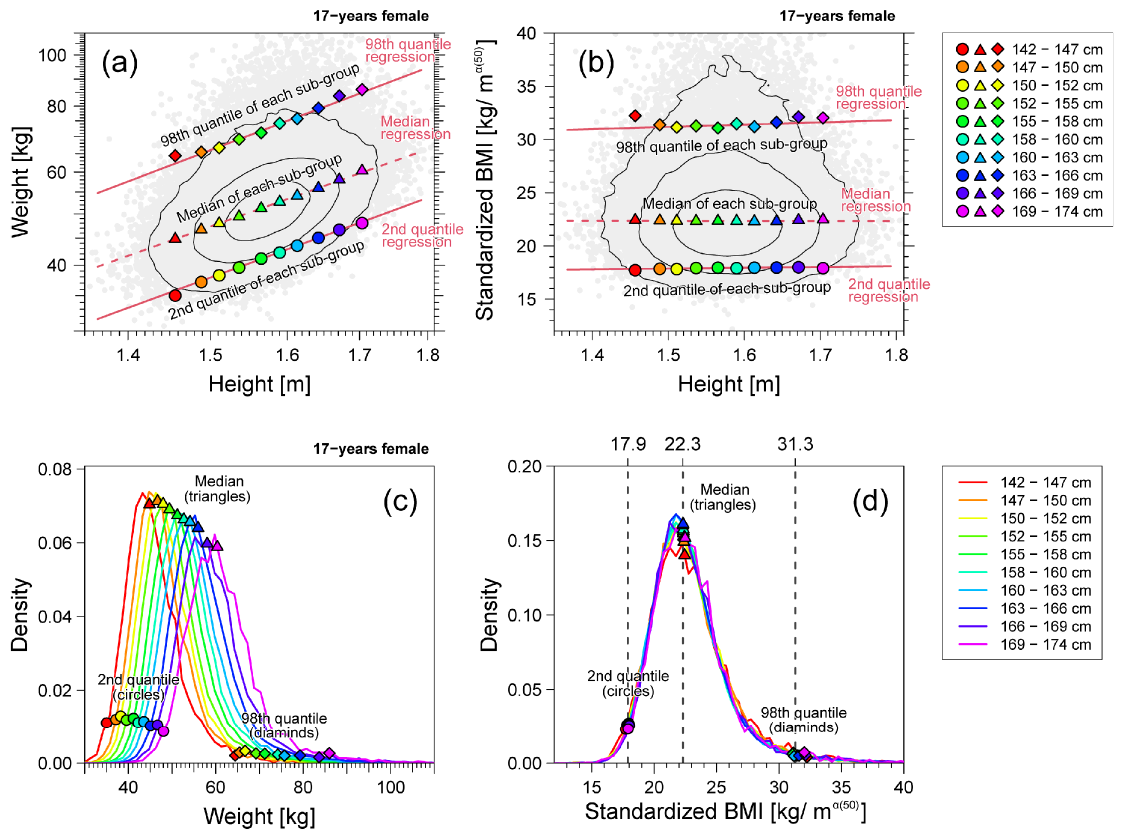
Weight and standardized BMI of 17-year-old females. Example of uni-scaling. (a) Weight-for-height distribution (gray points) in the log-log scale. (b) Standardized BMI *w/h*^1.85^-for-height distribution (gray points) in the linear scale. In panels (a) and (b), the contour lines are drawn with solid black lines; the 2nd, 50th, and 98th quantile regression curves are described by lines; the 2nd, 50th, and 98th centile points of the weight distribution in each height stratum are plotted as circles, triangles, and diamonds, respectively. (c) The weight distribution of each height stratum. (d) The standardized BMI *w/h*^1.85^ distributions of each height stratum. In panels (c) and (d), the 2nd, 50th, and 98th centile points of the distribution in each height stratum are plotted as circles, triangles, and diamonds, respectively. Ten height strata are defined in the range of the median ±3 SD.

In Fig. 9(b), quantile regression lines assuming a linear relationship between BMI and height are described with the estimated centile points of the BMI distribution in each height stratum. A horizontal regression line with an almost zero slope indicated that the BMI centile was almost independent of height. In this example, allometric uni-scaling further indicated the existence of an invariant BMI distribution, independent of height. The estimated distributions of BMI in the strata of different heights had almost identical shapes, as shown in Fig. 9 (d). However, as shown in Fig. 9 (c), the estimated weight distributions in the strata of different heights exhibit different shapes. This clearly shows the benefits of BMI as an excellent objective parameter for thinness and obesity assessment because the position of an arbitrarily selected centile was almost the same, independent of the height strata. The overall goal of developing a body build assessment index is to obtain the same index distribution at each height level [6]. In this sense, BMI is well-grounded and valid as a “body-build assessment index.”

Males usually stop growing and reach adult height by 16 or 17 years, whereas females do so by 14 or 15 years. Therefore, we expect BMI to be valid in young adults in a population-statistical sense as an excellent height-adjusted mass measure independent of height, age, and sex.

### Multi-scaling properties in children

Conventional BMI is not always an accurate objective body-build assessment measure because the scaling exponents *{α*(*q*)*}* in males and females under 17 years of age systematically deviate from 2 and exhibit multi-scaling properties, as shown in Fig. 8. Here, we explain how multi-scaling properties break down an objective comparison using a BMI-type formula. Figure 10 shows the same analysis as in Fig. 9 using 8-year-old female data. As shown in Fig. 10(a), the non-parallel regression lines demonstrate the multi-scaling property of the weight-for-height distribution. In this example, the estimated slope of the median regression line was *α*(50) = 2.73. Therefore, we modified the definition of BMI to use *w/h*^2.73^ as the standardized BMI. Figure 10(b) shows the relationship between the standardized BMI and height. Unlike Fig. 9(b), this result shows that the 2nd and 98th regression lines are no longer horizontal, which implies that the index distribution varies in a height-dependent manner. As shown in Fig. 10(d), the 2nd and 98th centile positions of the index are highly scattered, showing a solid height dependence in comparison with Fig. 9(d). This result indicates that no unified formula for *w/h*^*α*^, such as BMI, can be used to assess both thinness and obesity in children. Therefore, the use of different procedures to assess thinness and obesity is necessary. This will be discussed in the following section.

**Fig 10.**
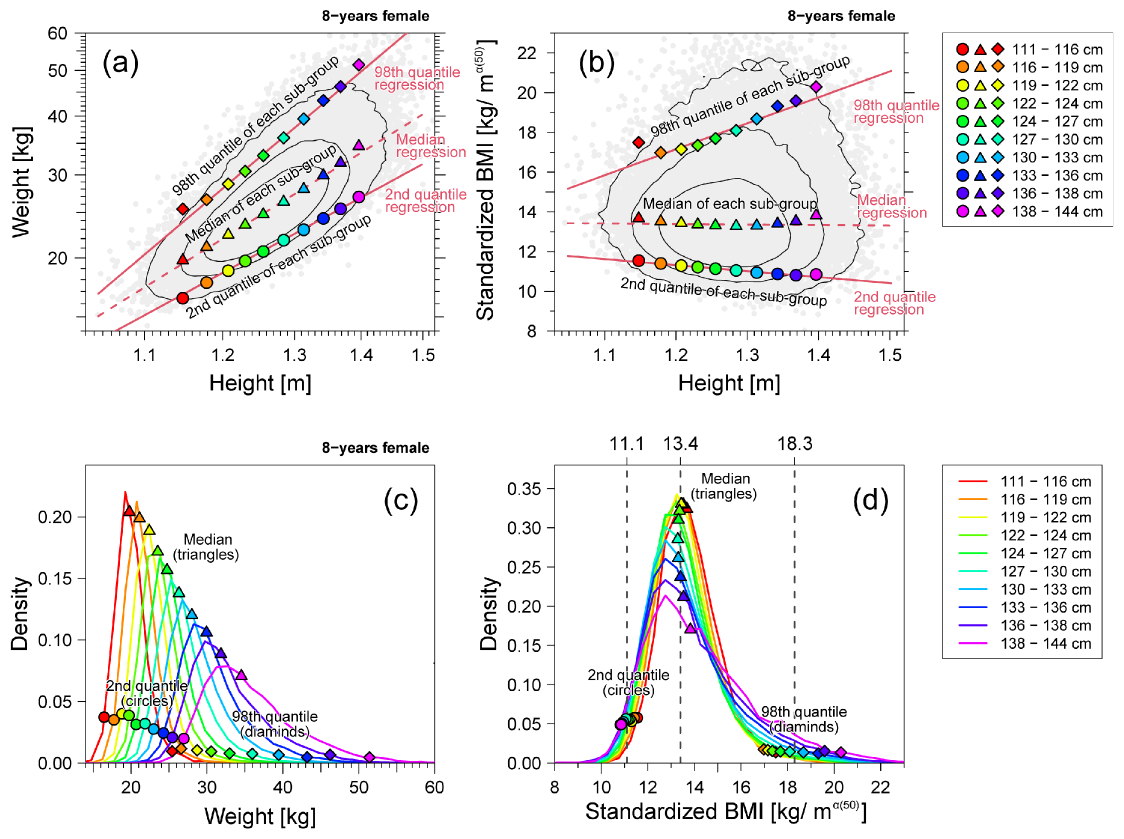
Weight and standardized BMI of 8-year-old females. Example of multi-scaling. (a) Weight-for-height distribution (gray points) in the log-log scale. (b) Standardized BMI *w/h*^2.73^-for-height distribution (gray points) in the linear scale. In panels (a) and (b), the contour lines are drawn with solid black lines; the 2nd, 50th, and 98th quantile regression curves are described by lines; the 2nd, 50th, and 98th centile points of the weight distribution in each height stratum are plotted as circles, triangles, and diamonds, respectively. (c) Weight distributions per height stratum. (d) Standardized BMI *w/h*^1.85^ distributions per height stratum. In panels (c) and (d), the 2nd, 50th, and 98th centile points of the distribution in each height stratum are plotted as circles, triangles, and diamonds, respectively. Ten height strata are defined in the range of the median ±3 SD.

Our finding of the multi-scaling property of the weight-for-height relationship in children indicates that the uncorrelated property of BMI and height assumed in previous studies is insufficient as a criterion for a height-adjusted mass measure. To explain this, we consider the schematic datasets shown in Fig. 11. We calculated the correlation coefficient between BMI and height, as shown in Figs. 11(a) and 11(b), both correlation coefficients were zero. However, in the example shown in Fig. 11(b), data with relatively large BMI and data with relatively small BMI in different height groups depend on the height. Therefore, when multi-scaling, BMI satisfying the uncorrelated criterion is not a height-adjusted measure in terms of population statistics.

**Fig 11.**
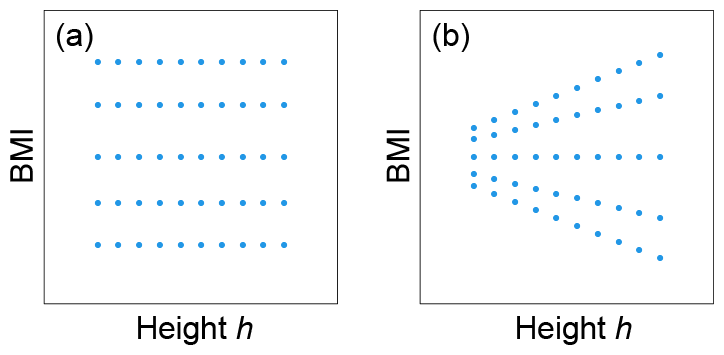
Limitation of uncorrelated criterion. Schematic depictions of the relationship between BMI and height. There is no correlation between BMI and height for either (a) or (b). Although panel (a) indicates no height dependence of each centile of BMI, panel (b) indicates height dependence of centiles except the median.

### Thinness and obesity assessment using extended BMI

In this study, we considered a method to overcome the shortcomings of the conventional BMI approach in assessing thinness and obesity in children and adolescents. To this end, we propose an extended BMI based on extended allometric scaling for the lower and upper centiles of the weight-for-height distribution for each sex and age group. The extended BMI can provide a height-adjusted measure for the assessment of thinness or fatness.

The extended BMI is defined as

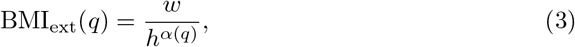

where *w* represents body weight in kilograms (kg), *h* represents height in meters (m), and *α* is the scaling exponent in Eq. (1), provided by the *q*th quantile regression of a reference dataset. When we used the extended BMI for the assessment of thinness, the cutoff value was provided by the model parameter *C*_*q*_ in Eq. (1).

Determining the cutoff values of an index for thinness and obesity is important. In the population-based approach, cutoff values were determined based on the centile points of the index distribution in the general population. To confirm this, we compare the BMI centile points in our data with the international cutoffs established by the International Obesity Task Force (IOFT) [38, 39]. Figure 12 plots the age dependence of the BMI centile points (solid lines) estimated from our data, together with the IOFT cutoff values (dashed lines). The results showed that the cutoff values for overweight were located in the 85th to 90th centiles, and the cutoff values for obesity were located in the 97th to 98th centiles. For thinness assessment, cut-off values for grade 1 thinness were located around the 10th centile, and cut-off values for grade 2 thinness were located at the 1st to 2nd centiles. The centile positions-vs-cutoff value relationship and the age dependence of the centile positions were almost identical for our data and other data from different countries [38, 39].

**Fig 12.**
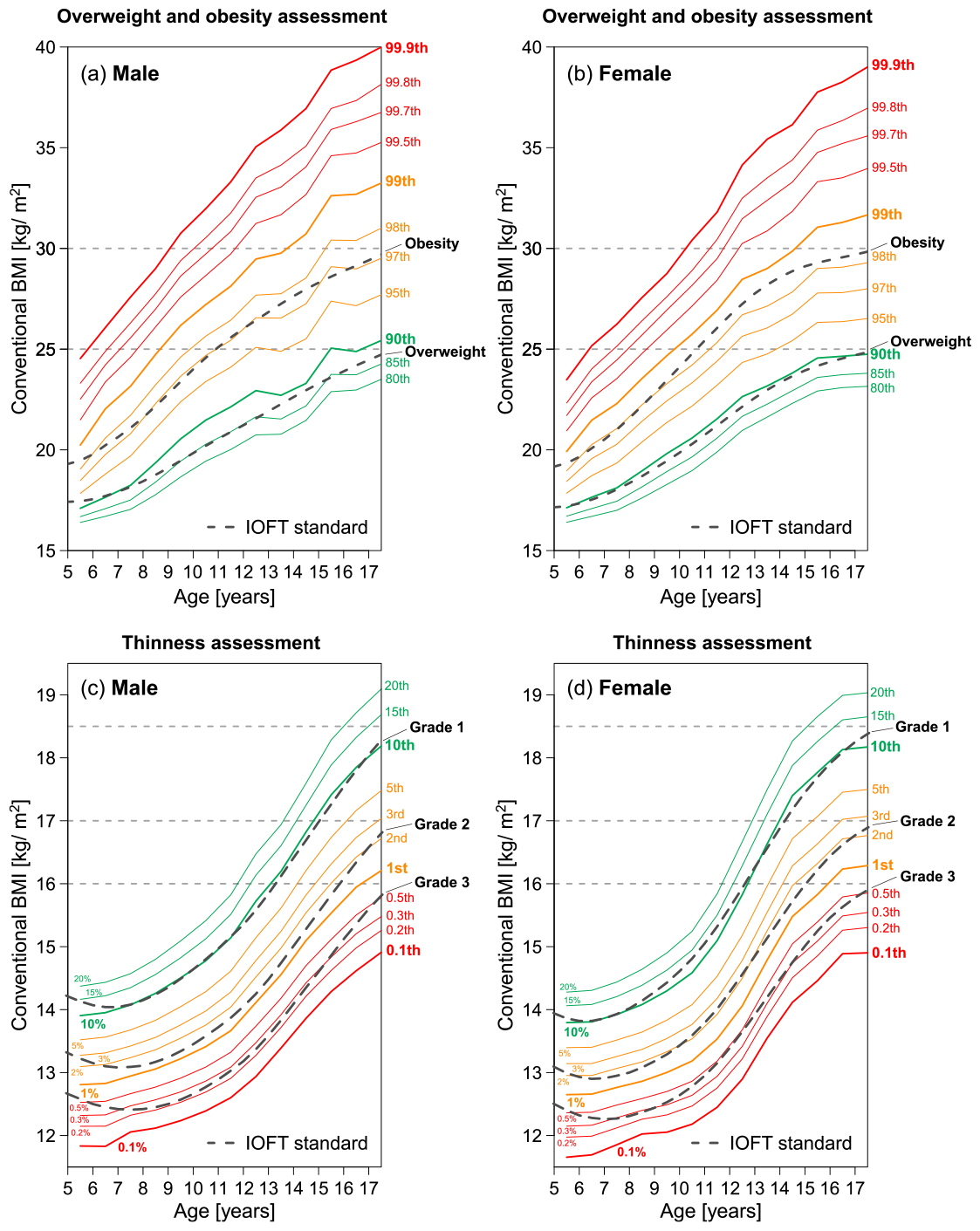
Comparison between BMI centiles and international cutoffs? The higher and lower centiles of our data are plotted with the international cutoffs established by the International Obesity Task Force (IOFT) [38, 39]. Age dependence of 80th to 99.9th centiles for male (a) and for female (b), and 0.1th to 20th centiles for male (c) and for female (d).

We propose an assessment method for thinness or fatness that uses an extended BMI instead of the conventional BMI. For instance, for the 2nd centile of the general population as the thinness cutoff, we use BMI_ext_(2) as the height-adjusted measure. This approach implies that the weights located at the 2nd centile in each weight group are matched among the different height groups, as depicted in the middle row of Fig. 13. In this case, BMI_ext_(2) cannot be employed for fatness assessment because BMI_ext_(2) for fatter individuals is widespread depending on height. Conversely, as indicated in the bottom row of Fig. 13, BMI_ext_(98) can be used to match the weights located at the 98th centile in each weight group among the different height groups, allowing for fatness assessment.

**Fig 13.**
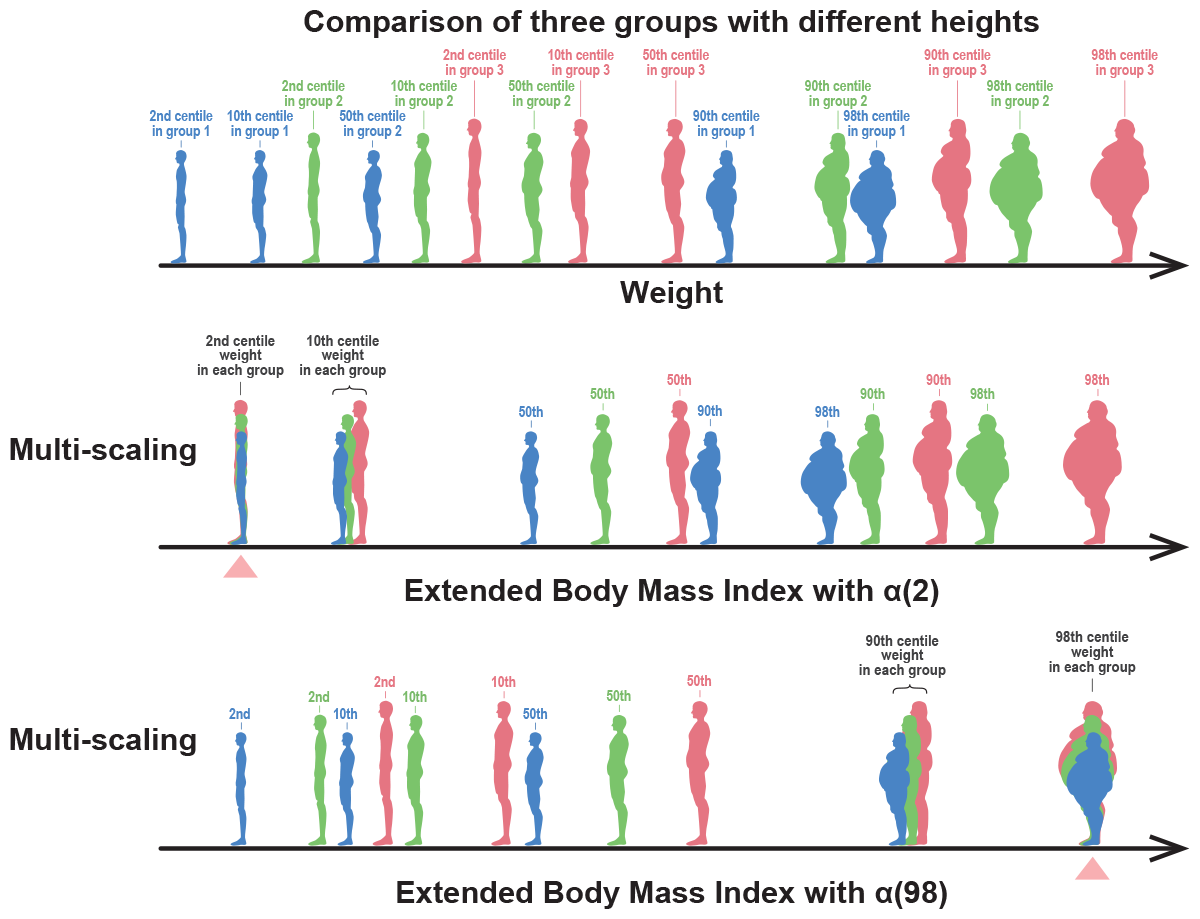
Meaning of extended BMI. The upper figure schematically depicts a body weight distribution of people in the 2nd, 10th, 50th, 90th, and 98th centile weights from the three different height groups (red, green, and blue represent the tall, medium, and short height groups, respectively). The middle figure depicts people shown in the upper figure sorted by their values of extended BMI (BMI_ext_(2)), in which the 2nd centile weight positions of each group coincide (indicated by triangle). The bottom figure depicts people shown in the upper figure sorted by their values of extended BMI (BMI_ext_(98)), in which the 98th centile weight positions of each group coincide.

Before explaining how the extended BMI works, we clearly show the drawbacks of conventional BMI in children and adolescents. Figure 14 shows the results for the 11-year-old males. The weight-for-height distributions shown in Fig. 14, the median and 2nd centile points of the weight are well approximated using Eq. (1) with scaling exponents of, respectively, 2.95 and 2.69 which are deviated from 2. The BMI-for-height distribution is shown in Fig. 14(b), the height dependence of the median and 2nd centile of BMI was observed. This height dependence implies that as shown in Fig. 14(d), the estimated distributions of BMI in strata of different heights shifted to the right as the height increased, and the median (triangles) and 2nd centile (circles) points of the BMI distributions were also shifted to the right. Therefore, BMI is not a height-adjusted measure for thinness assessment, because the locations of the 2nd centile points of BMI strongly depend on height. Our extended BMI can overcome this drawback of the conventional BMI.

**Fig 14.**
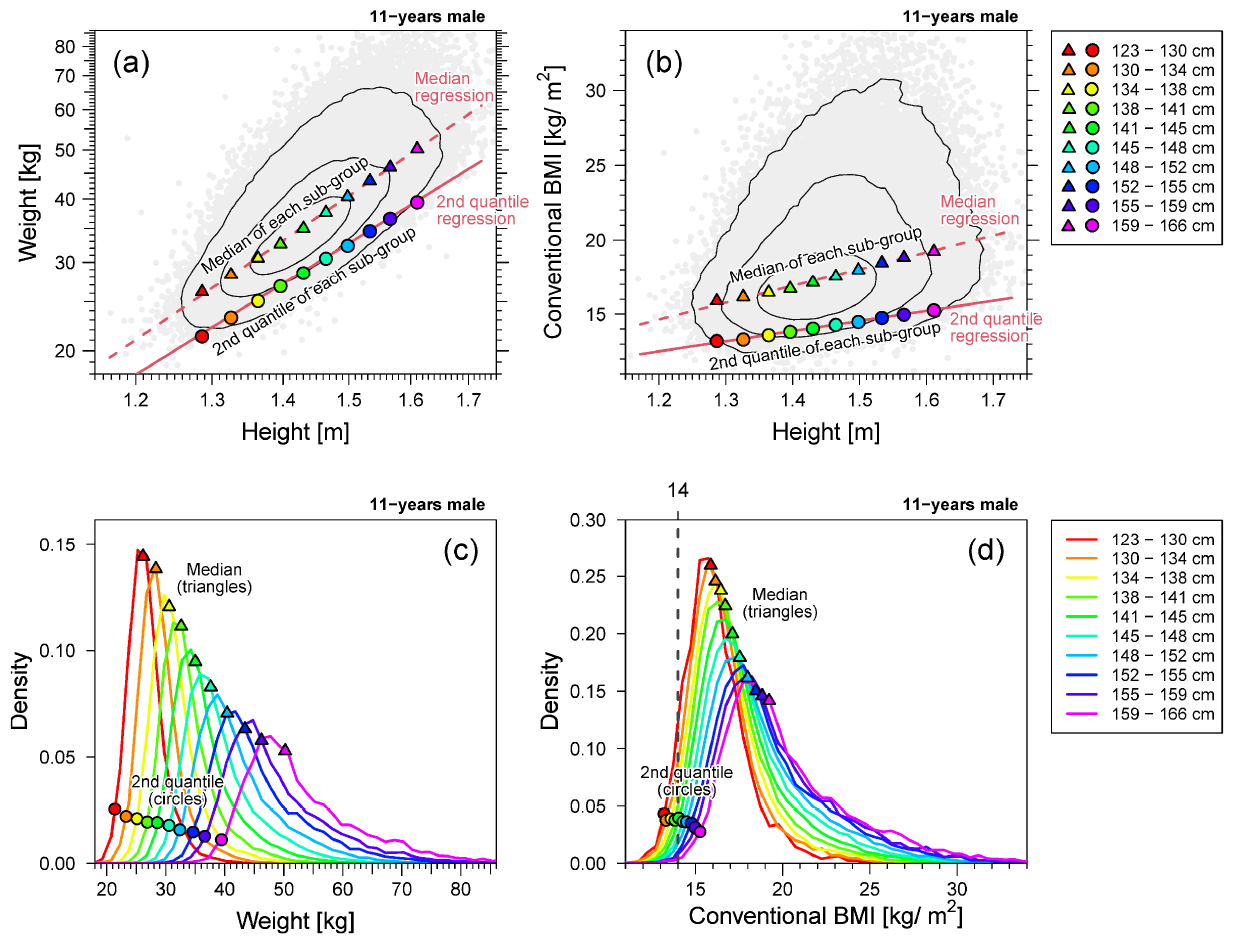
Weight and standardized BMI of 11-year-old males. Example of multi-scaling. (a) Weight-for-height distribution (gray points) in the log-log scale. (b) Conventional BMI *w/h*^2^-for-height distribution (gray points) in the linear scale. In panels (a) and (b), the contour lines are described by solid black lines; 2nd and 50th quantile regression curves are described by lines; 2nd and 50thcentile points of the weight distribution in each height stratum are plotted as circles and triangles, respectively. (c) Weight distributions of each height stratum. (d) Conventional BMI *w/h*^2^ distributions of each height stratum. In panels (c) and (d), the 2nd and 50th centile points of the distribution per height stratum are plotted as circles and triangles, respectively. Ten height strata are defined in the range of the median ±3 SD.

Here, we suppose the 2nd centile of weight as the cut-off for thinness assessment. Figure 15 shows the results when we define the extended BMI as BMI_ext_(2) = *w/h*^*α*(2)^ using the same data shown in Fig. 14. As shown in Fig. 15(a), the 2nd quantile regression line (solid line) of BMI_ext_(2) is horizontal, indicating that it is height-independent. In addition, the 2nd centile points (circles in Fig. 15(a)) of the BMI_ext_(2) distribution in the height strata are also horizontally aligned, although the 50th centile level is slightly slanted. As shown in Fig. 4 (b), the lower tails of the estimated distributions of BMI_ext_(2) in the strata of different heights almost overlap, demonstrating that BMI_ext_(2) is a good height-adjusted measure for thinness assessment.

**Fig 15.**
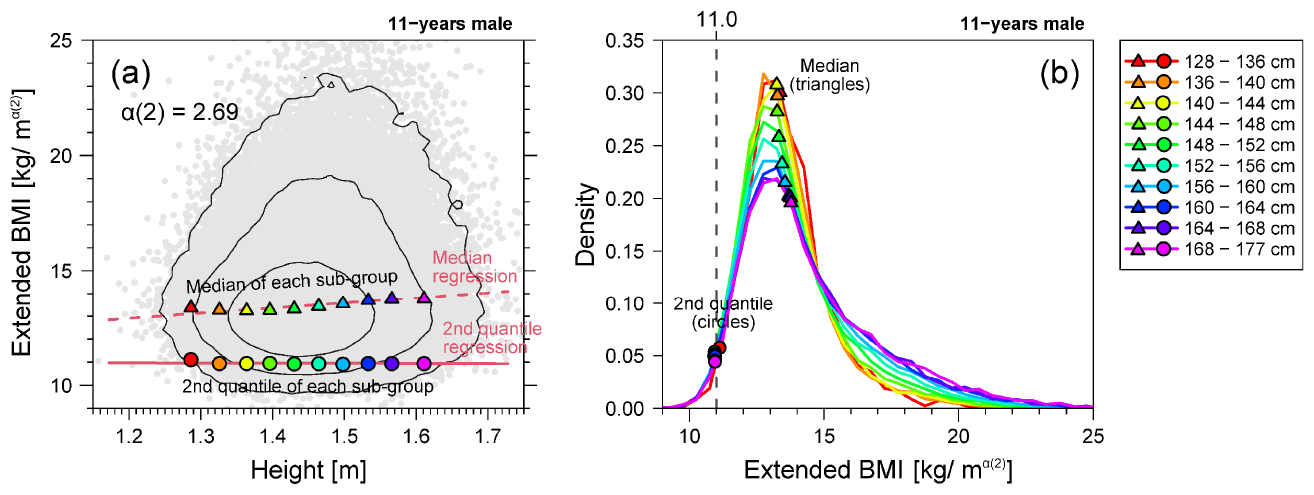
Extended BMI distributions of 11-year-old males. (a) BMI_ext_(2)-for-height distribution in the linear scale. (b) BMI_ext_(2) distributions of each height stratum. Figure formats of (a) and (b) are the same as Fig. 14(b) and (d), respectively, although for the extended BMI instead of conventional BMI.

Tables 2 and 3 summarize the scaling exponents and cutoff values for the thinness and obesity assessments. Unlike conventional BMI in adults, there were substantial age and sex dependences of the scaling exponents and cutoff values in children and adolescents. This, international standards for children and adolescents should be reviewed.

**Table 2.** Estimated parameters, *α*(*q*) and *C*_*q*_ for 2nd, 10th, 90th, and 98th centile for males.

| Age<br>[years] | 2nd centile |  | 10th centile |  | 90th centile |  | 98th centile |  |
| --- | --- | --- | --- | --- | --- | --- | --- | --- |
| | $\alpha(2)$ | $C_2$ | $\alpha(10)$ | $C_{10}$ | $\alpha(90)$ | $C_{90}$ | $\alpha(98)$ | $C_{98}$ |
| 5 | 2.06 | 13.02 | 2.13 | 13.73 | 2.61 | 16.12 | 3.17 | 16.87 |
| 6 | 2.21 | 12.73 | 2.30 | 13.34 | 3.10 | 14.93 | 3.72 | 15.45 |
| 7 | 2.34 | 12.41 | 2.40 | 13.02 | 3.45 | 13.59 | 3.98 | 14.00 |
| 8 | 2.39 | 12.19 | 2.49 | 12.68 | 3.76 | 12.40 | 4.05 | 13.41 |
| 9 | 2.49 | 11.84 | 2.57 | 12.36 | 3.95 | 11.50 | 4.08 | 12.95 |
| 10 | 2.57 | 11.48 | 2.67 | 11.93 | 3.92 | 11.24 | 3.91 | 13.18 |
| 11 | 2.69 | 11.00 | 2.73 | 11.64 | 3.59 | 12.07 | 3.63 | 14.06 |
| 12 | 2.68 | 11.05 | 2.71 | 11.77 | 3.23 | 13.54 | 3.27 | 15.93 |
| 13 | 2.70 | 10.98 | 2.72 | 11.68 | 3.16 | 13.13 | 3.24 | 15.34 |
| 14 | 2.60 | 11.58 | 2.61 | 12.46 | 2.93 | 14.58 | 3.00 | 17.18 |
| 15 | 2.36 | 13.33 | 2.34 | 14.60 | 2.43 | 20.02 | 2.51 | 23.35 |
| 16 | 2.17 | 15.02 | 2.13 | 16.71 | 2.10 | 23.56 | 2.26 | 26.46 |
| 17 | 2.06 | 16.16 | 2.02 | 18.03 | 2.02 | 25.19 | 2.13 | 28.90 |

**Table 3.** Estimated parameters, *α*(*q*) and *C*_*q*_ for 2nd, 10th, 90th, and 98th centile for females.

| Age<br>[years] | 2nd centile |  | 10th centile |  | 90th centile |  | 98th centile |  |
| --- | --- | --- | --- | --- | --- | --- | --- | --- |
| | $\alpha(2)$ | $C_2$ | $\alpha(10)$ | $C_{10}$ | $\alpha(90)$ | $C_{90}$ | $\alpha(98)$ | $C_{98}$ |
| 5 | 2.04 | 12.91 | 2.12 | 13.65 | 2.54 | 16.33 | 2.94 | 17.30 |
| 6 | 2.19 | 12.62 | 2.28 | 13.28 | 3.01 | 15.25 | 3.47 | 16.05 |
| 7 | 2.32 | 12.32 | 2.38 | 12.96 | 3.22 | 14.24 | 3.64 | 14.90 |
| 8 | 2.38 | 12.07 | 2.47 | 12.63 | 3.39 | 13.45 | 3.70 | 14.30 |
| 9 | 2.50 | 11.62 | 2.58 | 12.15 | 3.41 | 13.08 | 3.60 | 14.28 |
| 10 | 2.67 | 10.93 | 2.74 | 11.48 | 3.34 | 12.95 | 3.45 | 14.53 |
| 11 | 2.81 | 10.37 | 2.87 | 10.93 | 3.33 | 12.80 | 3.41 | 14.52 |
| 12 | 2.74 | 10.79 | 2.77 | 11.56 | 2.91 | 15.42 | 3.12 | 16.61 |
| 13 | 2.50 | 12.28 | 2.42 | 13.85 | 2.28 | 20.54 | 2.54 | 21.42 |
| 14 | 2.15 | 14.96 | 2.02 | 17.22 | 1.97 | 24.14 | 2.29 | 24.51 |
| 15 | 1.87 | 17.33 | 1.84 | 19.11 | 1.89 | 25.75 | 2.14 | 27.21 |
| 16 | 1.89 | 17.56 | 1.88 | 19.17 | 1.83 | 26.63 | 2.07 | 28.18 |
| 17 | 1.92 | 17.38 | 1.91 | 18.94 | 1.79 | 27.17 | 1.95 | 29.96 |

### Interpretation of allometric multi-scaling

Finally, we discuss the interpretation of multi-scaling and emphasize that the scaling exponents estimated by our extended allometric analysis provide interesting and significant insights into growth process dynamics in children and adolescents.

Mathematically, the allometric scaling relation *w* = *Ch*^*α*^ can be described as a solution to the differential equation

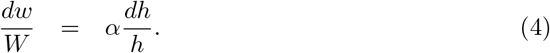

In this equation, *dw/w* (weight increase divided by weight) represents the body weight gain rate and *dh/h* (height increase divided by height) represents the body height growth rate . Therefore, the scaling exponent quantifies the ratio between the body weight gain rate and body height increase rate:

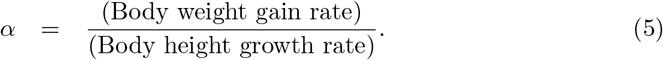

In the following, we assume that the value of *α* is related to the body weight gain rate along the time axis of the height growth process for children.

Multi-scaling properties of the weight-for-height distribution in children and adolescents imply that the growth balance of height and weight strongly depends on the weight centile position in each height group. This centile dependence means that the body weight gain rate of relatively overweight children in the same age and height groups is remarkably greater than the average (or median) weight of the children. This tendency is more pronounced at ages under 15 years for males and under 12 years for females, as shown in Fig. 8, overweight children at these ages were more likely to accelerate their weight gain than average-weight children. Conversely, the body weight gain rate of relatively underweight children was lower than that of the average weighted children under 12 years of age for both males and females. Thus, being underweight at these ages could be more likely to decelerate weight gain and cause growth retardation. Such growth balance differences, depending on thinness and fatness, disappear at 17 years of age, close to adulthood, in both males and females. To effectively manage the nutritional status of children, the origin and adverse effects of growth balance differences between height and weight, depending on thinness and fatness, must be clarified. Determining whether the origin lies in genetic or social factors related to diet will help to identify effective obesity prevention measures for children.

From a biophysical viewpoint, weight-for-height scaling, *w ∝ h*^2^, can be linked to Kleiber’s law, which states that resting energy expenditure (metabolic rate) is proportional to *w*^3*/*4^ [29]. In addition to Kleiber’s law, the “surface law” is also known, asserting that if species of different sizes maintain the same body temperature, their heat production should be related to the surface area, not body mass [28, 40]. Based on this law, we assume that human energy expenditure is proportional to the body surface area *S*. The combination of Kleiber’s law and the surface law implies that

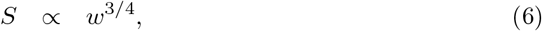

where *∝* denotes the proportional relationship.

Furthermore, if we approximate the shape of the human body using cylindrical parts, the surface area *S* is given by a function of *w* and *h* (see details in Ref. [29, 41]).

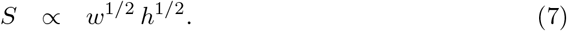

Values of the scaling exponents can be interpreted as the weight increase rate adjusted for height.

Using Eqs. (6) and (7), we get

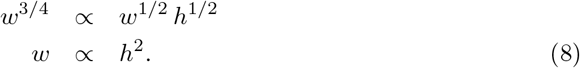

This result is consistent with that of the conventional BMI formula for adults. Note that the validity of Kleiber’s and surface laws is still under debate, and the above discussion is a rough estimation, but is empirically reasonable.

Our findings indicated that these considerations are not applicable to children. The metabolic rate in children is known to be significantly higher than that predicted by Kleiber’s law [42], and it has been suggested that a higher metabolic rate in children is related to the relative growth rate of the internal organs that contribute largely to the resting metabolic rate [40]. For example, five internal organs (the brain, heart, kidneys, liver, and lungs) occupy 14.6% of the total body mass of a 10 kg child but only 6.3% of the total body mass of a 70 kg adult [42]. Furthermore, the brain accounts for 9.2% of the total body weight and 45% of the total metabolic rate in children weighing 10 kg, compared with 2% of the total body weight and 21% of the total metabolic rate in adults weighing 70 kg. In addition, differences in body shape between children and adults may cause multi-scaling in children. Rapid changes in body composition balance may be associated with multi-scaling in children and adolescents.

The above discussion suggests that the age-dependence of multi-scaling properties could be closely related to growth process dynamics. Particularly, for female, as illustrated in Fig. 16, age-dependent characteristic changes in the scaling exponents of different could be associated with major developmental phenomena in the female growth process, such as the adolescent growth spurt (peak height age), menarche, and height growth stop (final height age). As females grow, all scaling exponents of different centiles begin to decrease around the age at which the adolescent growth spurt occurs and then begin to change to uni-scaling after the age of menarche (Fig. 16). At ages immediately after height growth stops, the indices asymptotically approach uni-scaling, with the scaling exponent *α*(*q*) = 2. Furthermore, if we regard the scaling exponent *α*(*q*) as a surrogate measure for the weight gain rate, our analysis results suggest that being underweight in females under 12 years of age results in growth retardation compared to average-weight children. As depicted in Fig. 16, the decreasing period of α(2) from 12 to 15 years old shows about one year delay compared with that of *α*(50). Because the weight gain rate approximated by α(2) for underweight females after age 12 is greater than *α*(50) for average-weight females, this phenomenon may imply a compensated recovery from delayed weight gain. We refer to this phenomenon as “growth retardation recoupment.” Thus, extremely low weight in females under 12 years of age may be associated with severe irreparable health issues.

**Fig 16.**
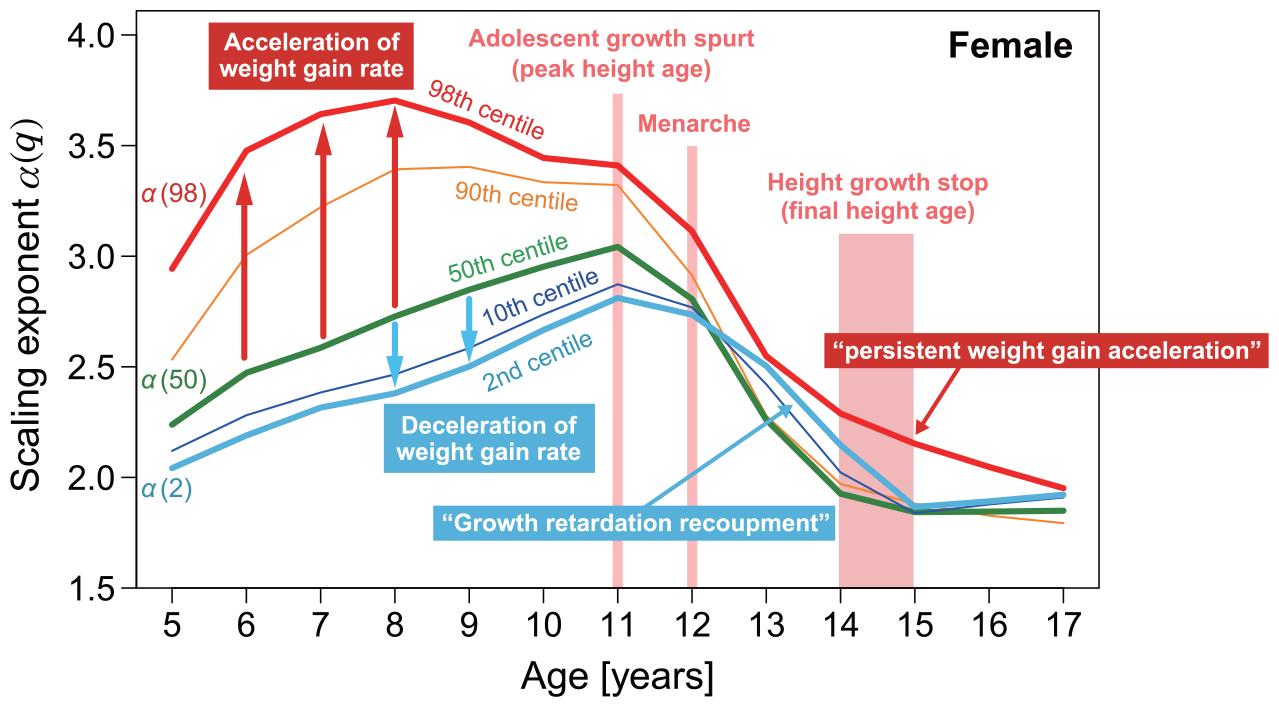
Age dependence of scaling exponents. The average ages at which the major developmental phenomena occur are indicated on the same plot as Fig. 8.

By contrast, in overweight females characterized by the 98th centile curve (red solid line in Fig. 16), the weight gain rate approximated by *α*(98) remained persistently higher than that of lighter-weight females. This suggests that the persistent tendency for accelerated weight gain in children under the age of 17 years is likely to induce more severe obesity. We refer to this phenomenon as “persistent weight gain acceleration.” Our data suggest that in females younger than 12 years of age, underweight causes growth retardation, and overweight causes persistent weight gain acceleration. This study analyzed only Japanese data, which is a limitation of this study. In the future, an extensive international database should be analyzed to understand these developmental changes and establish new criteria for assessing thinness and obesity in children and adolescents.

## Conclusion

In this study, we proposed a mathematical condition called uni-scaling, in which an objective height-adjusted measure for thinness and obesity assessment must be fulfilled from the viewpoint of population statistics. The population statistical validity of a height-adjusted measure means that people with different heights can be compared using weight-centile positions within the same height group. To ensure the validity of a height-adjusted measure, the uncorrelated criterion used in previous studies to test the correlation between a measure and height is insufficient because lower and higher distributions can have height dependence. By introducing an extended allometric analysis estimating the scaling exponent of any given centile of the weight-for-height distribution, and by analyzing a large-scale Japanese database including children aged 5-17 years, we showed the importance of the uni-scaling criterion and the existence of multi-scaling in children and adolescents.

Our results demonstrated that the conventional BMI for 17 years in both males and females fulfilled the uni-scaling criterion and had the same scaling exponents of 2, independent of the weight-for-height centile levels. Therefore, BMI of 17-year-old males, females and young adults is a well-grounded and valid height-adjusted index for thinness and obesity assessment in terms of population statistics. We also demonstrated remarkable multi-scaling properties at ages 5-13 years for males and 5-11 years for females, and convergence to uni-scaling with a scaling exponent close to 2 as they approached 17 years of age for both sexes. Our results confirmed that conventional BMI is not population-statistically valid for thinness and obesity assessment in males under 16 years and females under 14. Instead of the conventional BMI, we propose an extended BMI to assess thinness and obesity in children and adolescents.

As summarized in Fig. 1, the issues of BMI validity have been discussed in terms of the validity of the formula [33] and the cutoff [38, 39]. In contrast, we focused on the objectivity of BMI or the meaning of height adjustment, and proposed a method to verify it. This is the decisive difference between our study and the previous studies. Although we demonstrated the difficulty in assessing children’s thinness and fatness by analyzing simple data on age, sex, height, and weight, such simple data can provide significant insights into the growth process dynamics of children and adolescents using extended allometric analysis. Our analysis results offer important hypotheses: being underweight in females under 12 years of age causes growth retardation recoupment, and being overweight in females under 17 years of age induces persistent weight gain acceleration. In the future, it will be necessary to establish a better understanding and assessment methods for thinness and fatness in children and adolescents. To this end, in addition to the population statistical approach, it is necessary to accumulate findings from a wide range of perspectives, including physiological, epidemiological, genetic, and sociological.

